# The bone microenvironment invigorates metastatic seeds for further dissemination

**DOI:** 10.1101/2020.11.16.383828

**Authors:** Weijie Zhang, Igor L. Bado, Hai Wang, Swarnima Singh, Hin-Ching Lo, Muchun Niu, Zhan Xu, Jun Liu, Weiyu Jiang, Yi Li, Jingyuan Hu, Xiang H.-F. Zhang

## Abstract

Metastasis has been considered as the terminal step of tumor progression. However, recent clinical studies suggest that many metastases are seeded from other metastases, rather than primary tumors. Thus, some metastases can further spread, but the corresponding pre-clinical models are lacking. By using several approaches including parabiosis and an evolving barcode system, we demonstrated that the bone microenvironment facilitates breast and prostate cancer cells to further metastasize and establish multi-organ secondary metastases. Importantly, dissemination from the bone microenvironment appears to be more aggressive compared to that from mammary tumors and lung metastases. We further uncovered that this metastasis-promoting effect is independent from genetic selection, as single cell-derived cancer cell populations (SCPs) exhibited enhanced metastasis capacity after being extracted from the bone microenvironment. Taken together, our work revealed a previously unappreciated effect of the bone microenvironment on metastasis evolution, and suggested a stable reprogramming process that engenders cancer cells more metastatic.

## INTRODUCTION

Metastasis to distant organs is the major cause of cancer-related deaths. Bone is the most frequent destination of metastasis in breast cancer and prostate cancer (Gundem et al., 2015; Kennecke et al., 2010; Smid et al., 2008). In the advanced stage, bone metastasis is driven by the paracrine crosstalk among cancer cells, osteoblasts, and osteoclasts, which together constitute an osteolytic vicious cycle (Esposito et al., 2018; Kang et al., 2003; Kingsley et al., 2007; Weilbaecher et al., 2011). Specifically, cancer cells secrete molecules such as PTHrP, which act on osteoblasts to modulate the expression of genes including RANKL and OPG (Boyce et al., 1999; Juárez and Guise, 2011). The alterations of these factors in turn boost osteoclast maturation and accelerate bone resorption. Many growth factors (e.g., IGF1) deposited in the bone matrix are then released, and reciprocally stimulate tumor growth. This knowledge laid the foundation for clinical management of bone metastases (Coleman et al., 2008).

The urgency of bone metastasis research is somewhat controversial. It has long been noticed that, at the terminal stage, breast cancer patients usually die of metastases in multiple organs. In fact, compared to metastases in other organs, bone metastases are relatively easier to manage. Patients with the skeleton as the only site of metastasis usually have better prognosis than those with visceral organs affected (Coleman and Rubens, 1987; Coleman et al., 1998). These facts argue that perhaps metastases in more vital organs should be prioritized in research. However, metastases usually do not occur synchronously. In 45% of metastatic breast cancer cases, bone is the first organ that shows signs of metastasis, much more frequently compared to the lungs (19%), liver (5%) and brain (2%) (Coleman and Rubens, 1987). More importantly, in more than two-thirds of cases, metastases will not be limited to the skeleton, but rather subsequently occur to other organs and eventually cause death (Coleman, 2006; Coleman and Rubens, 1987; Coleman et al., 1998). This raises the possibility of secondary dissemination from the initial bone lesions to other sites. Indeed, recent genomic analyses concluded that the majority of metastases result from seeding from other metastases, rather than primary tumors (Brown et al., 2017; Gundem et al., 2015; Ullah et al., 2018). Thus, it is imperative to investigate further metastatic seeding from bone lesions, as it might lead to prevention of the terminal stage, multi-organ metastases that ultimately cause the vast majority of deaths.

Despite its potential clinical relevance, little is known about metastasis-to-metastasis seeding. Current preclinical models focus on seeding from primary tumors, but cannot distinguish between additional sites of dissemination. We have recently developed an approach, termed intra-iliac artery injection (IIA), that selectively deliver cancer cells to hind limb bones via the external iliac artery (Wang et al., 2015, 2018; Yu et al., 2016). Although it skips the early steps of the metastasis cascade, it focuses the initial seeding of tumor cells in the hind limbs, and allows the tracking of secondary metastases from bone to other organs. It is therefore a suitable model to investigate the clinical and biological roles played by bone lesions in multi-organ metastasis-to-metastasis seeding.

## RESULTS

### Temporally lagged multi-organ metastases in mice carrying IIA-introduced bone lesions of breast and prostate cancers

IIA injection has been employed to investigate early-stage bone colonization. Both aggressive (e.g., MDA-MB-231) and relatively indolent (e.g. MCF-7) breast cancer cells can colonize bones albeit following different kinetics. In both cases, cancer cell distribution is highly bone-specific at early time points, allowing us to dissect cancer-bone interactions without the confounding effects of tumor burden in other organs (Figure 1A) (Wang et al., 2015, 2018). However, as bone lesions progress, metastases (as indicated by bioluminescence signals) begin to appear in other organs, including additional bones, lungs, liver, kidney, and brain, usually 4-8 weeks after IIA injection of MDA-MB-231 cells (Figure 1B, 1C and S1). This phenomenon is not specific for the highly invasive MDA-MB-231 cells, but was also observed in more indolent MCF-7 cells and PC3 prostate cancer cells, albeit after a longer lag period for PC3 cells (8-12 weeks) (Figure 1D-G). Remarkably, in many cases, counter-lateral hind limbs (designated as “L.Hindlimb” for “left hind limb” as the initial bone lesions were introduced to the right hind limb) are affected in all models examined.

**Figure 1.**
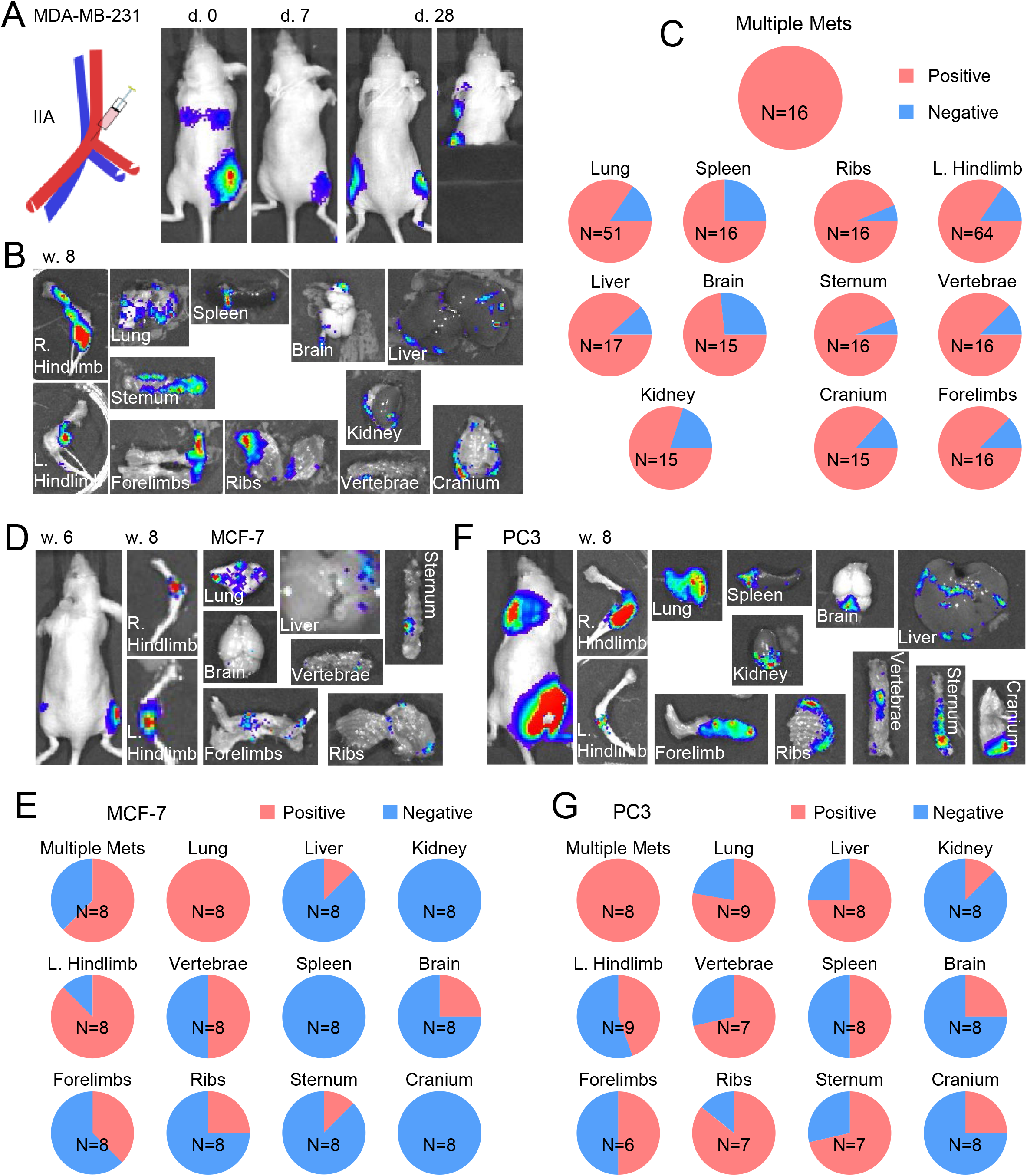
Temporally lagged multi-organ metastases in mice carrying bone lesions introduced by intra-iliac artery (IIA) injection. **(A)** Left, diagram of intra iliac artery (IIA) injection. Right, representative bioluminescence images showing the distribution pattern of tumor cells in different time points. 1E5 MDA-MB-231 fLuc-mRFP (MDA-MB-231 FR) triple negative breast cancer cells were injected through iliac artery to induce bone lesions at the right hindlimb of 6-week old female nude mice. **(B)** Representative *ex vivo* bioluminescent images show the metastatic spread across multiple tissues in nude mice with MDA-MB-231 FR cells inoculated in bone after 8 weeks. **(C)** Frequencies of metastatic involvement were quantified as determined by *ex vivo* bioluminescent imaging. Hereafter, the threshold for positive involvement was set as 15 photon/pixel with 120 seconds exposure time. The frequency of ‘multiple metastases’ represents the frequency of nude mice which show the presence of metastases at least 3 tissues other than the primary site (IIA, right hindlimb; IIV or TV, lung; MFP or MIND, mammary gland). The total number of mice examined were indicated. **(D)** Representative *in vivo* and *ex vivo* bioluminescent images show the development of multiple site metastases in nude mice receiving 1E5 non-metastatic MCF-7 luminal breast cancer cells through IIA injection. **(E)** Frequencies of multiple sites metastases in nude mice receiving MCF-7 cells through IIA were quantified by *ex vivo* bioluminescent imaging of various tissues. **(F)** Representative *in vivo* and *ex vivo* bioluminescent images show the development of multiple site metastases in male nude mice receiving 2E5 PC3 prostate cancer cells through IIA injection. **(G)** Frequencies of multiple sites metastases in nude mice receiving PC-3 cells through IIA were quantified by *ex vivo* bioluminescent imaging of various tissues.

### Intra-iliac vein (IIV) injection as a control/ comparison demonstrated enhanced metastatic capacity of cancer cells in bone lesions

The later-appearing multi-organ metastases may result from further dissemination of cancer cells in the initial bone lesions. Alternatively, they could also arise from cancer cells that leaked and escaped from bone capillaries during IIA injection. In the latter case, the leaked cancer cells would enter the iliac vein and subsequently arrive in the lung capillaries. Indeed, there did appear to be bioluminescence signals in lungs upon IIA injection (Figure 1A). To distinguish these probabilities, we performed IIV injection with the same cell quantity, to be compared with the results of IIA injection at late time points. The IIV injection procedure should mimic the “leakage” from IIA injection, although allowing many more cells to enter the venous system (Figure 2A, compared to Figure 1A). Strikingly, animal subjected to IIV injection of MDA-MB-231 cells developed much fewer metastases to almost every organ examined except for lungs (Figure 2B-D, S2A, and S2B). These results strongly favor the secondary metastasis hypothesis and indicate that pre-existing bone lesions are associated with much higher metastatic burdens in other organs. Although the lungs are an exception, it should be noted that IIV injection allows many more cells to enter the pulmonary circulation compared to IIA injection (Figure S2C). Yet, the two approaches still resulted in similar lung metastatic burdens, suggesting that tumor cells disseminated from bone lesions have higher lung metastasis efficiencies than cells directly injected into venous circulation (Figure S2D). Similar results were obtained from the more indolent MCF-7 cells, in which case even lung signals showed a significant difference between IIA and IIV groups (Figure 2E and S2E).

**Figure 2.**
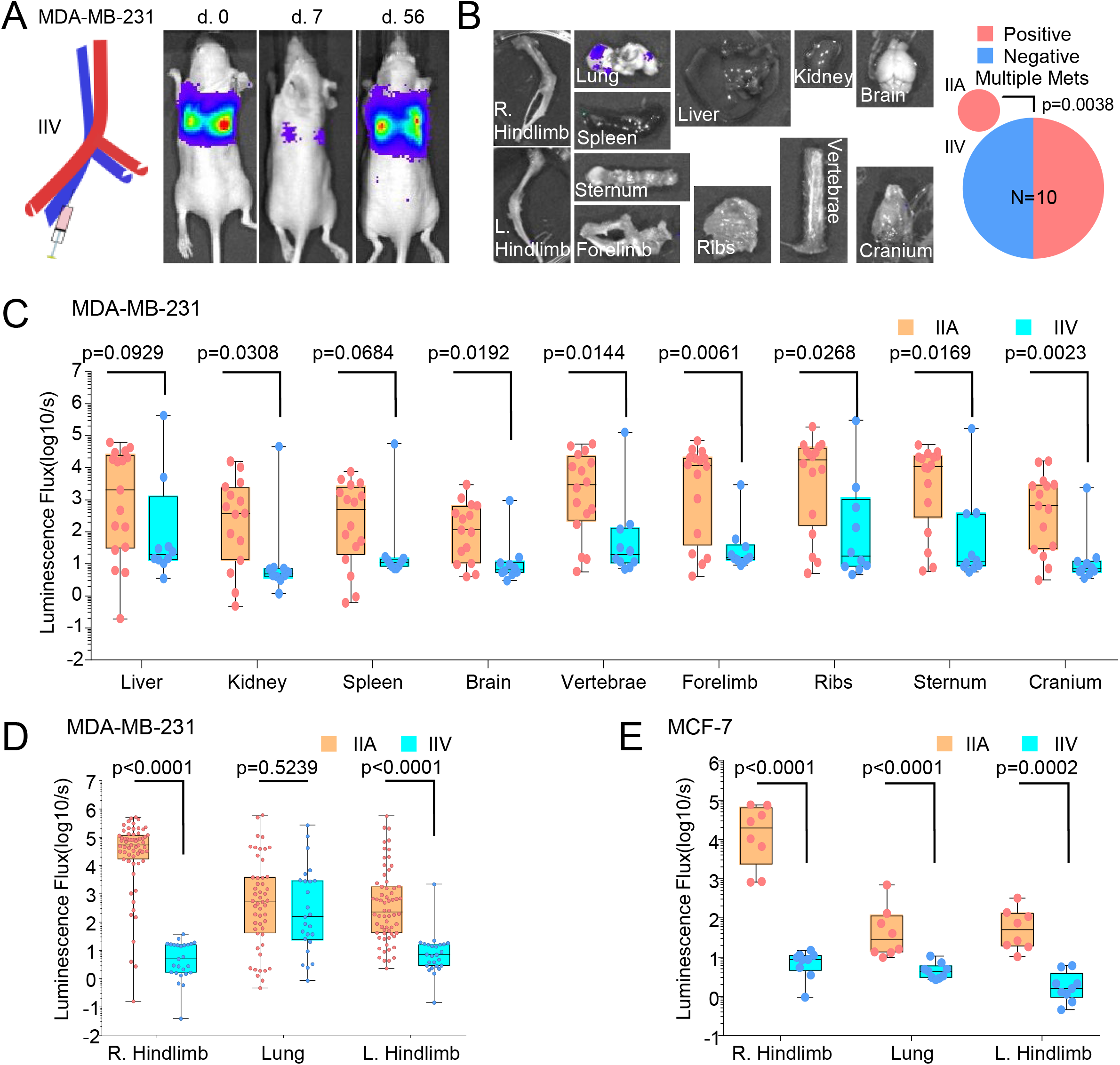
Intra-iliac vein (IIV) injection, as compared to IIA injection, results in much lower frequency of multi-organ metastases. **(A)** Left, diagram of intra-iliac vein injection (IIV). Right, representative bioluminescence images at indicated time points. 1E5 MDA-MB-231 FR cells were injected through iliac vein to bypass the bone marrow vasculature of the right hindlimb in 6-week old female nude mice. **(B)** Representative *ex vivo* bioluminescent images (Left) and quantification (Right) of the incidence of multi-organ metastases in mice receiving the same number of MDA-MB-231 cells through IIV injection as compared to those of IIA injected ones. P value was determined by Fisher’s exact test. **(C)** Combined boxplot and dot plots quantitating the metastatic burden in indicated tissues in mice receiving MDA-MB-231 cells through IIA injection compared to IIV injection, P values were determined by Mann-Whitney test. **(D)** The same as (C), but focusing on lungs and counter-lateral hind limb (L.Hindlimb). **(E)** Combined boxplot and dot plots quantitating the metastatic burden in indicated tissues in mice receiving MCF-7 cells through IIA injection compared to IIV injection, P values were determined by Mann-Whitney test.

### Bone provides a unique environment that promotes further metastases

We asked if the metastatic seeding from bone lesions is more efficient as compared to that from orthotopic tumors and metastatic lesions in other organs. To this end, we first performed mammary fat pad (MFP) injection to introduce orthotopic tumors (Figure 3A), and assessed metastatic distribution at the same time points as with previous experiments. Spontaneous metastases from mammary tumors were detected in multiple organs as would be expected for MDA-MB-231 tumors (Figure 3B). However, compared to IIA-injected mice, MFP-injected mice showed significantly lower frequencies and signal intensities of secondary metastatic lesions in the vast majority of organs (Figure 3B-D, and S3A). This cannot be explained by different tumor loads in the source tumors: the mammary tumors in MFP-injected mice exhibited much higher bioluminescence signal intensity compared to that of right hind limb bone lesions in IIA-injected mice (Figure S3B). The exception is again the lungs, where a similar metastatic burden was observed (Figure 3C), which highlights the multi-organ nature of secondary metastases from bones, in contrast to lung-tropic metastases from orthotopic tumors, at least in the MDA-MB-231 model.

**Figure 3.**
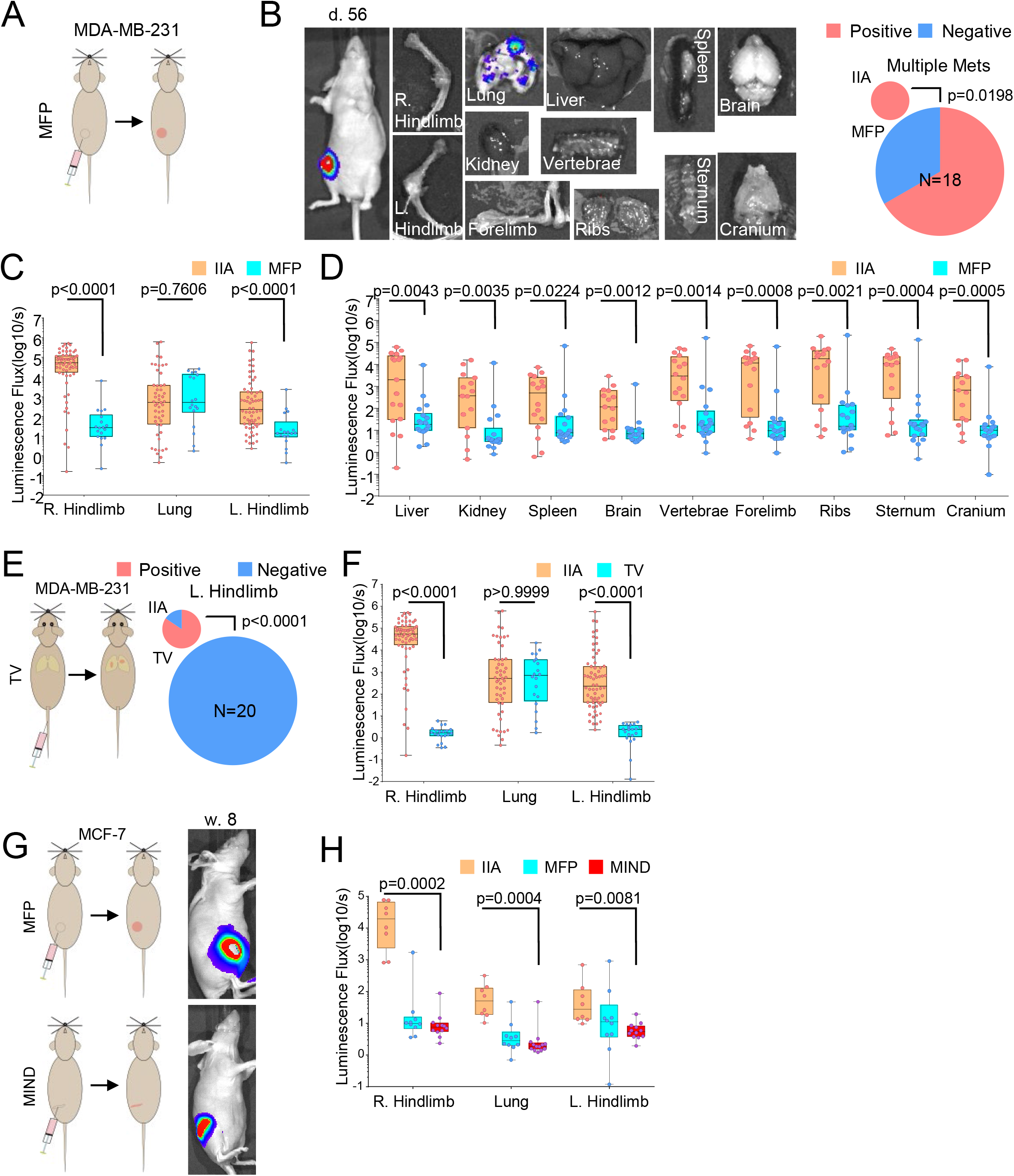
The specificity of bone microenvironment in promoting further metastasis. **(A)** Diagram showing the mammary fat pad implantation (MFP) of 1E5 MDA-MB-231 FR cells in nude mice. **(B)** Representative *ex vivo* bioluminescent images (Left) and quantification (Right) show that less multi--organ metastases occur in mice receiving the same number of MDA-MB-231 cells through MFP injection as compared to IIA injected ones. **(C)** Combined boxplot and dot plots quantitating the metastatic burden in indicated tissues in mice receiving MDA-MB-231 FR cells through IIA injection compared to MFP injection, P values were determined by Mann-Whitney test. **(D)** The same as (C) but in a wider range of soft tissues and bones. **(E)** Left: schematic shows the comparison group subjected to tail vein (TV) injection. Right: Pie charts show the proportion of mice that develop multi-organ metastases in IIA-injected mice versus TV-injected mice, Fisher’s exact test was used to determine the P value. **(F)** Combined boxplot and dot plots quantitating the metastatic burden in indicated tissues in mice receiving MDA-MB-231 cells through IIA injection compared to TV injection. P values were determined by Mann-Whitney test. **(G)** Schematic shows comparison groups that receive either MFP or intra ductal implantation (MIND) injection. **(H)** Combined boxplot and dot plots quantitating the metastatic burden in indicated tissues in mice receiving MCF-7 cells through MFP injection and MIND injection. P values were determined by Kruskal-Wallis test.

We next carried out tail-vein (TV) injection to examine secondary metastases from established lung lesions. Interestingly, very few secondary metastases were observed (Figure 3E and 3F), suggesting that the lung microenvironment does not facilitate further dissemination.

We then turned to the more indolent MCF-7 cells, which are luminal-like and express estrogen receptor (ER). MCF-7 was traditionally considered as a “non-metastatic” model. Recently, it was shown that mammary intra-ductal injection of MCF-7 cells could result in multi-organ metastases (Sflomos et al., 2016). We thus conducted both MFP and intra-ductal injection (referred to as “MIND” model) with the same cell numbers, and compared the capacity of multi-organ metastases (Figure 3G). When the injected tumor lesions reach similar sizes as measured by bioluminescence intensity (Figure S3C), significantly more metastatic lesions were found in the IIA group compared to the other two groups (Figure 3H and S3D). Taken together, these results suggested that cancer cells in the bone exhibited enhanced ability to further metastasize to other organs.

### Cross-seeding of cancer cells from bone lesions to orthotopic tumors

Cancer cells may enter circulation and seed other tumor lesions or re-seed the original tumors (Kim et al., 2009). By using MDA-MB-231 cells tagged with different fluorescent proteins, we asked if bone lesions can cross-seed mammary tumors (Figure 4A). Interestingly, we observed that while orthotopic tumors can be readily seeded by cells derived from bone lesions, the reverse seeding did not seem to occur (Figure 4B and 4C). This difference again highlights the enhanced metastatic aggressiveness of cancer cells in the bone microenvironment.

**Figure 4.**
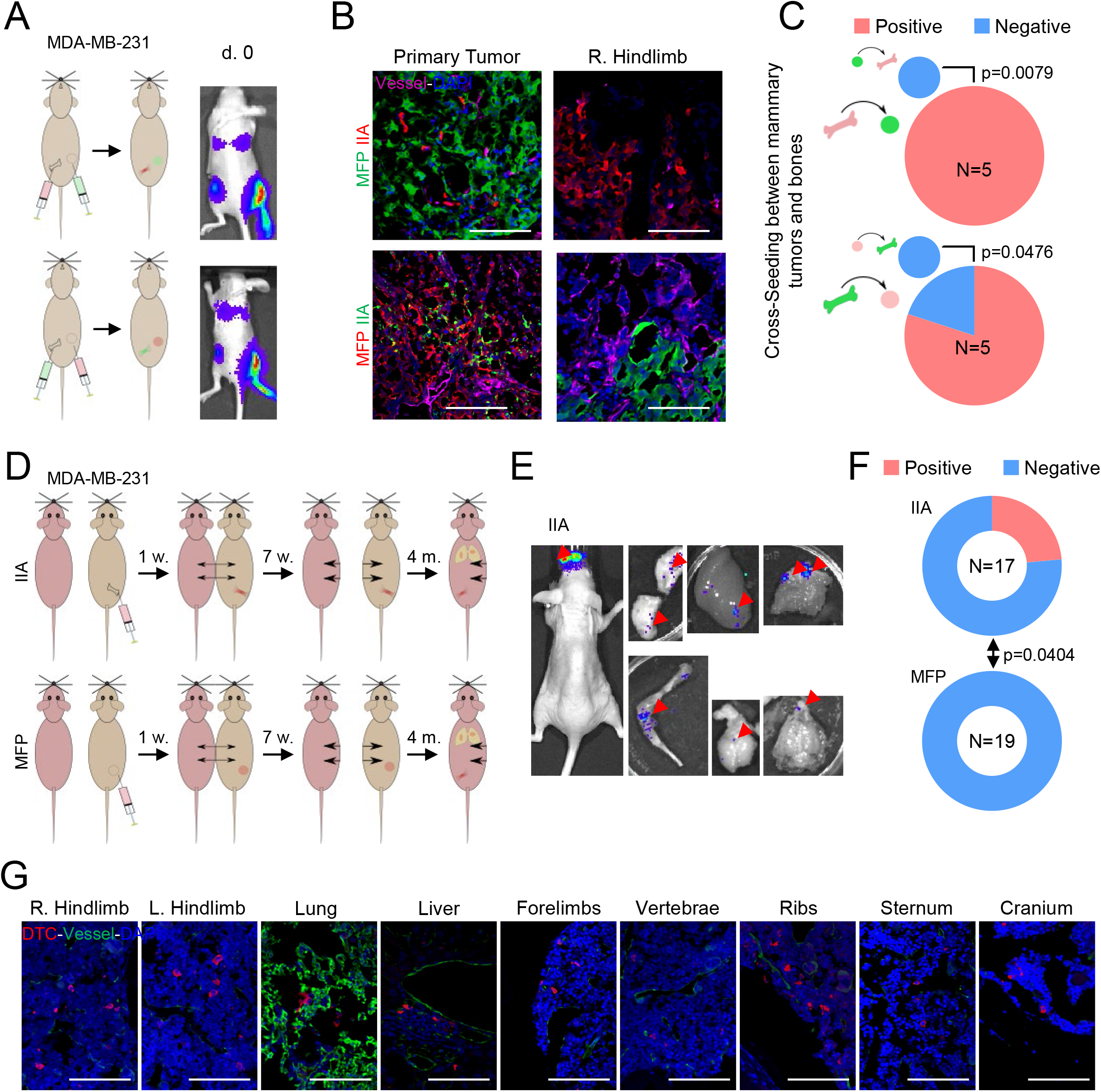
Cross-seeding and parabiosis experiments support the promoting effects of bone microenvironment in further metastases. **(A)** Schematics show the experimental design of cross-seeding experiment between primary tumors and bone lesions. Upper, IIA injection with mRFP-tagged MDA-MB-231 cells and MFP implantation with EGFP-tagged MDA-MB-231 cells; lower, the colors were swapped between bone lesions and mammary tumors. **(B)** Representative confocal images show that the seeding from bone metastases to primary tumors and vice versus. Blood vessels are also stained (magenta). **(C)** Incidence of cross-seeding between bone lesions and mammary tumors in both directions. P value was determined by Fisher’s exact test. **(D)** Schematics show the experimental design to compare the dissemination capacity of bone metastases and primary tumors using parabiosis model. Briefly, nude mice were implanted with 1E5 MDA-MB-231 cells via IIA (upper) or MFP (lower) 1 week prior to the parabiosis surgery. The parabiotic pairs were maintained for 7 weeks to allow the fusion of circulation between tumor-bearing donor mice and recipient tumor-free mice, before a surgical separation was performed. The recipient mice were then continuously monitored by weekly bioluminescent imaging until metastatic signals were detected in up-to 4 months. **(E)** Representative bioluminescent images showing that the metastatic disease occurs in parabiotic mice from IIA group. **(F)** Parabiotic mice from IIA group are prone to develop metastases, whereas those from MFP group did not form metastases. P value was determined by Fisher’s exact test. **(G)** Representative confocal images confirm the presence of tumor cells in various tissues of parabiotic mice of IIA group.

### Parabiosis models support enhanced capacity of cancer cells to metastasize from bone to other organs

It is possible that IIA injection disturbs bone marrow and stimulates systemic effects that allow multi-organ metastases. For example, the injection might cause a transient efflux of bone marrow cells that can arrive at the distant organ to form pre-metastatic niche. To test this possibility, we used parabiosis to fuse the circulation between a bone lesion-carrying mouse (donor) and tumor-free mouse (recipient) one week after IIA injection. In parallel, we also performed parabiosis on donors that have received MFP injection and tumor-free recipients (Figure 4D). After seven weeks, surgical separation was performed to allow time for metastasis development in the recipients. Subsequently, the organs of originally tumor-free recipients were collected and examined for metastases four months later. Only ~20% of recipients in the IIA group were found to harbor cancer cells in various organs (Figure 4E and 4F), mostly as microscopic disseminated tumor cells (Figure 4G), indicating that the fusion of circulation system is not efficient for metastatic seeds to cross over from donor to recipient. However, in the MFP comparison group, no metastatic cells were detected (Figure 4F, S4A, and S4B), and th difference is statistically significant. Therefore, the parabiosis data also support the hypothesis that the bone microenvironment invigorates further metastasis, and this effect is unlikely to be due to IIA injection-related systemic influence.

### An evolving barcode system revealed the phylogenetic relationships between initial bone lesions and secondary metastases

Barcoding has become widely used to elucidate clonal evolution in tumor progression and therapies. An evolving barcoding system has recently been invented for multiple parallel lineage tracing (Kalhor et al., 2017, 2018). It is based on CRISPR/Cas9 system, but utilizes guide RNAs that are adjacent to specific protospacer adjacent motif (PAM) in their genomic locus, thereby allowing Cas9 to mutate its own guide RNAs. These variant guide RNAs are named homing guide RNAs (hgRNAs). When Cas9 is inducibly expressed, hgRNA sequences will randomly drift as a function of time, serving as evolving barcodes (Figure 5A, and S5A). Therefore, over time, individual clones of cells will accumulate more and more different mutations to their barcodes, allowing the identification of distinct lineages.

**Figure 5.**
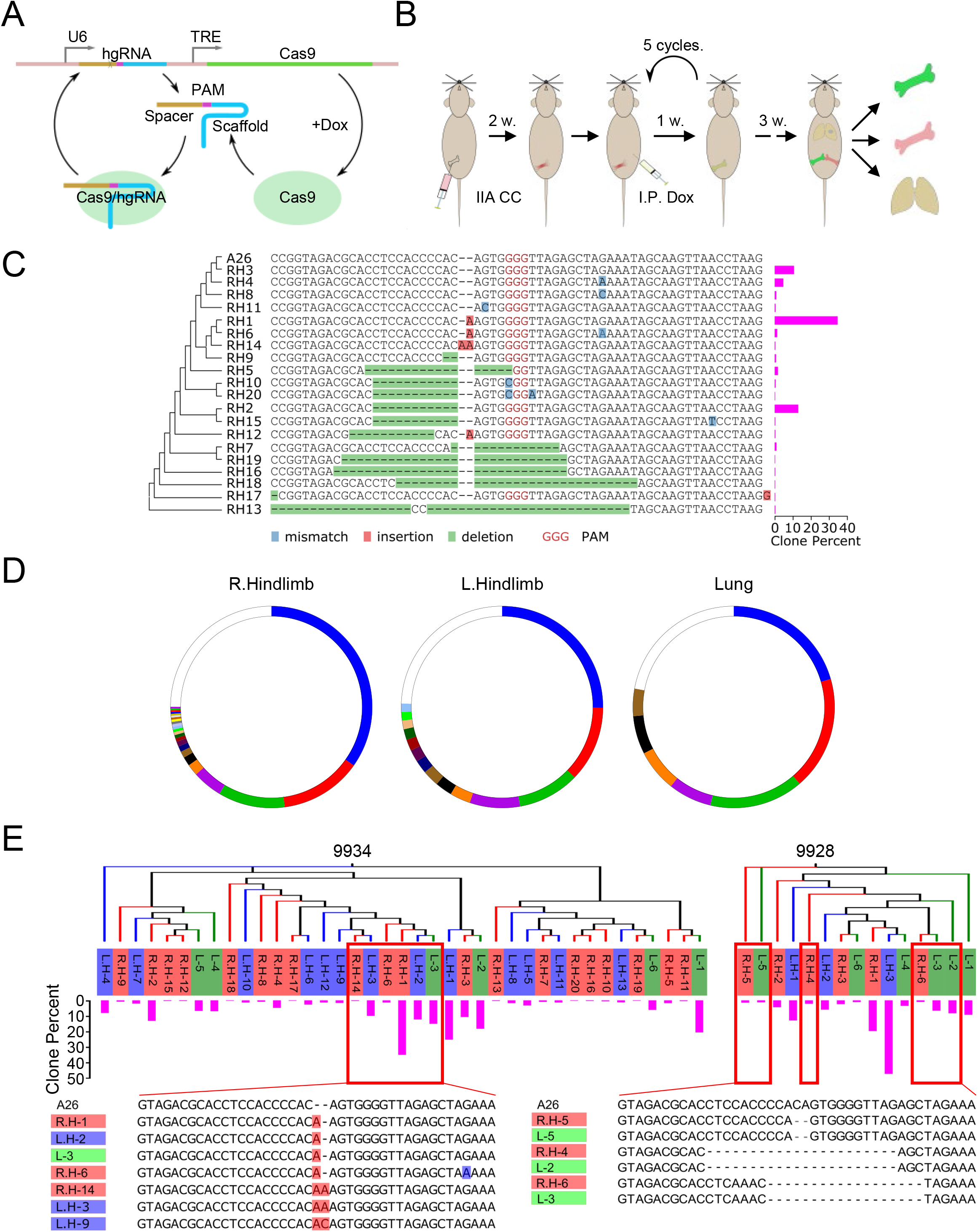
Metastatic evolution delineated by an evolving barcode system. **(A)** A schematic shows the principle of the evolving barcode system comprised of homing guide RNAs (hgRNAs) and inducible Crispr-Cas9. The PAM sequence was inserted into the spacer and scaffold sequence to allow the self-targeting of Cas9-hgRNA complex. Dox treatment induces the expression of Cas9 protein, and thereby introduce mutations on the spacer sequences (Kalhor et al., 2017, 2018). **(B)** A schematic shows the experimental design using the evolving barcoding system to study the metastatic spread *in vivo*. MDA-MB-231 FR cells were transduced with TLCV2-hgRNA A26 and selected by puromycin for 2 weeks. 1E5 barcoded cells were then delivered to bone via IIA. 2 weeks later, a single dose of 5mg/kg doxycycline weekly for 5 cycles was applied to the mice carrying bone metastases through I.P. injection. 3 weeks after Dox treatment, lung, right and left hindlimbs were collected, and total DNA were exacted. The barcodes regions were PCR amplified, and sequenced with Illumina Next-Seq. **(C)** Representative images showing the multiple sequence alignment of mutated barcodes extracted from metastases in indicated organs. **(D)** Representative images showing the clonal composition of metastases at right hindlimb, left hind limb and lung from mouse 9934. **(E)** Neighbor-Joining phylogenetic trees without distance corrections suggest the majority of dominant clones from left hindlimbs and lungs are closely related to the clones from bone lesions at right hindlimbs.

We transduced MDA-MB-231 cells using a barcode with a relatively slow mutation rate (Figure S5B). Upon IIA injection, doxycycline was used to induce expression of Cas9, and hence the onset of barcode evolution. When secondary metastases developed, we harvested them as well as the initial bone lesions and performed barcode sequencing (Figure 5B). A wide spectrum of mutations to the barcodes was found (Figure 5C), allowing us to trace the evolution of multiple clones. Furthermore, we observed lower clonal diversity (Figure 5D) and clear phylogenetic connections between the dominant clones in secondary metastases and major clones in initial bone lesions (Figure 5E and S5C). Overall, these results are consistent with our expectation that the majority of secondary metastases are derived from the initial bone lesions, rather than cells that leaked during the injection (Figure S5D). Interestingly, many secondary metastasis lesions contain cells derived from multiple clones in the bone lesions, which suggest that there are multiple seeding events occurring either sequentially or simultaneously. This is consistent with recent genomic studies showing that metastases are mostly multi-clonal (Siegel et al., 2018).

### The enhanced secondary metastatic seeding from bone involves a process independent of clonal selection

Organo-tropism is an important feature of metastasis. Clonal selection appears to play an important role in organ-specific metastasis, which has been intensively studied previously (Bos et al., 2009; Kang et al., 2003; Minn et al., 2005; Vanharanta and Massague, 2013). Herein, the metastasis-promoting effects of the bone microenvironment appear to be multi-organ and do not show specific organ-tropism. In an accompanied study, we discovered profound phenotypic shift of ER+ breast cancer cells in the bone microenvironment, which included loss of luminal features and gain of stem cell-like properties (Bado et al., 2019). This shift is expected to promote further metastases (Gupta et al., 2019; Ye and Weinberg, 2015). Therefore, we hypothesize that the enhancement of metastasis may be partly through an epigenomic dedifferentiation process. To test this possibility, we compared the metastasis capacity of a genetically identical SCP of MCF-7 cells and its bone-entrained derivative (the same cell line extracted from bone lesions). We used intra-cardiac injection to simultaneously deliver cancer cells to multiple organs (Figure 6A). As expected, bone-entrained SCP2 was more capable of colonizing distant organs and gave rise to much higher tumor burden in multiple sites (Figure 6B-D). qRT-PCR revealed that the bone-entrained SCP2 overexpressed multiple cancer stem cell markers and mesenchymal markers while maintaining the expression of epithelial markers (Figure 6E-F), which is consistent with the result of reverse phase protein array (RPPA) in the accompanied study (Bado et al., 2019). Altogether, this suggests that exposure to the bone environment alters tumor cell gene expression and molecular characteristics such as stem cell properties and epithelial phenotype, which ultimately promotes their metastatic ability (Li and Kang, 2016).

**Figure 6.**
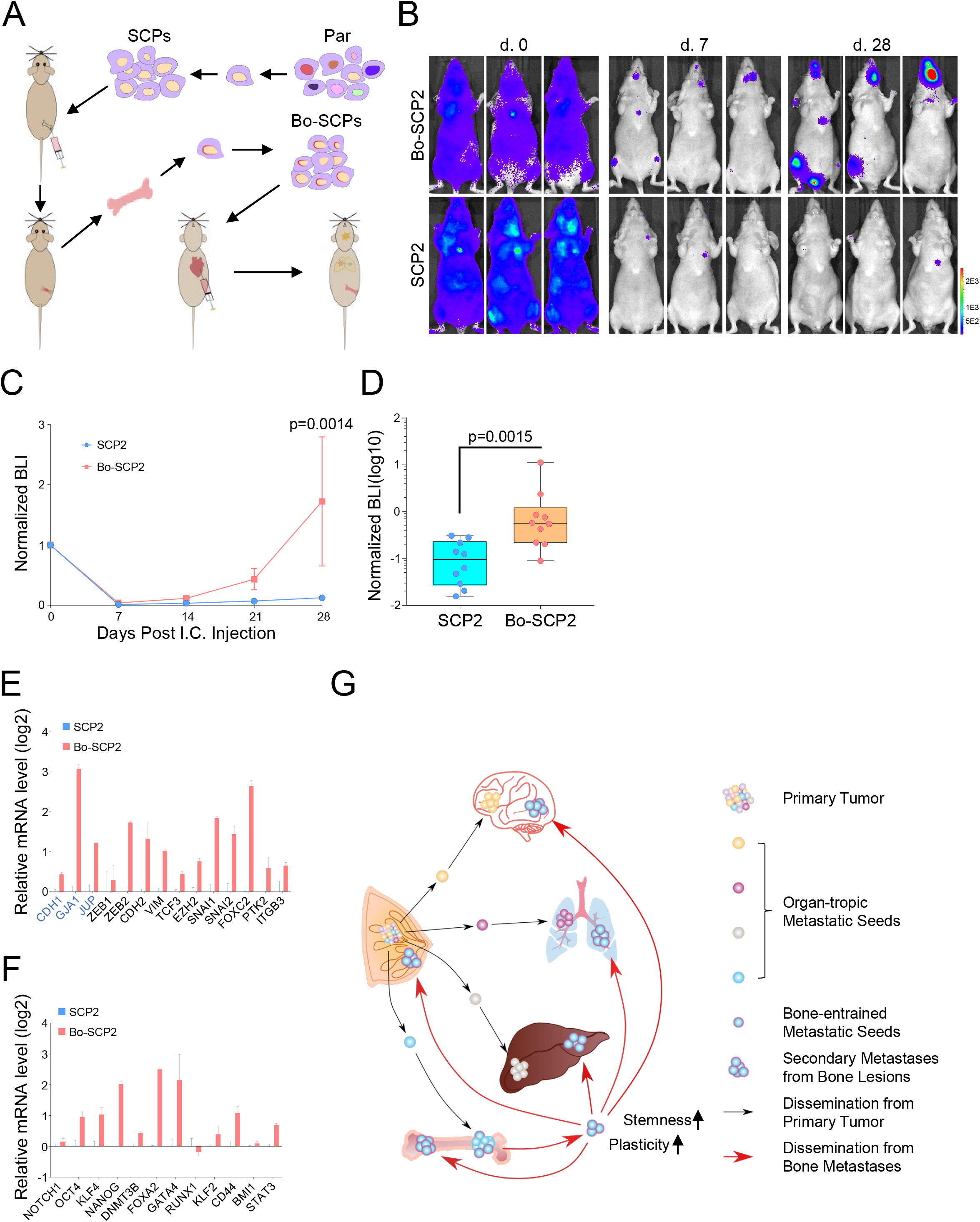
Bone-entraining boosts metastatic capacity of single cell-derived cancer cells. **(A)** Experimental design to test the metastatic capacity of bone-entrained single cell derived population (SCP). MCF-7 SCP2 cells were inoculated in bone for 6 weeks, extracted, and expanded in vitro. The metastatic capacity of bone-entrained SCP2 (SCP2-Bo) were examined by intra-cardiac (I.C.) injection. **(B)** Representative bioluminescent images show the colonization kinetic of SCP2-Bo and parental SCP2 cells. **(C)** The growth kinetics of SCP2-Bo and parental SCP2 cells *in vivo* after intra-cardiac injection. LSD test was used to determine the p value. **(D)** The comparison of the normalized increase between SCP2-Bo and parental SCP2 cells at the endpoint, p values were determined by Mann-Whitney test. **(E)** Relative mRNA levels of EMT markers in SCP2-Bo and parental SCP2 cells. Blue, Epithelial markers; Black, Mesenchymal markers or EMT promoters. **(F)** Relative mRNA levels of stemness markers in SCP2-Bo and parental SCP2 cells. **(G)** Models for secondary metastatic dissemination from existing bone metastases. The organ-tropic metastatic seeds are pre-existing in the primary tumor and may constitute the first wave of dissemination from the primary tumor. The bone tropic seeds are epigenetically reprogrammed by the bone microenvironment, which leads to the enhanced plasticity and stemness. These bone-entrained metastatic seeds exhibit reduced organo-tropism but increased metastatic capacity, which may constitute the secondary wave of dissemination from bone lesions.

## DISCUSSION

In this study, based on the IIA injection technique and through multiple independent approaches, we demonstrated that the bone microenvironment not only permits cancer cells to further disseminate but also appears to augment this process. A key question that remains is the timing of secondary metastasis spread out of the initial bone lesions: whether this occurs before or after the bone lesions become symptomatic and clinically detectable. The answer will determine if therapeutic interventions should be implemented in adjuvant or metastatic settings, respectively. Moreover, if further seeding occurs before bone lesions become overt, it raises the possibility that metastases in other organs might arise from asymptomatic bone metastases, which might warrant further investigations. Indeed, our co-submitted study indicated that in the early phase of bone colonization, cancer cells already acquire stem cell-like features (Bado et al., 2019), supporting that asympotomatic bone micrometastases are potentially capable of metastasizing before being diagnosed. Future studies will be needed to precisely determine the onset of secondary metastasis from bones.

The fact that the genetically homogenous SCP cells became more metastatic after lodging into the bone microenvironment suggest a mechanism distinct from genetic selection. Remarkably, this phenotype persists even after in vitro expansion, so it is relative stable and suggests an epigenomic reprogramming process, which has been characterized in depth in the accompanied study (Bado et al., 2019). We propose that this epigenetic mechanism may act in concerted with the genetic selection process. Specifically, the organ-specific metastatic traits may pre-exist in cancer cell populations (Minn et al., 2005; Zhang et al., 2013), and determine the first site of metastatic seeding. The epigenomic alterations will then occur once interactions with specific microenvironment niches are established and when cancer cells become exposed chronically to the foreign milieu of distant organs. Our data suggest that such alterations drive a second wave of metastases in a less organ-specific manner (Figure 6G). This may explain why terminal stage of breast cancer is often associated with multiple metastases (DiSibio and French, 2008).

In the clinic, some bone metastases can be managed for years without further progression, while others quickly develop therapeutic resistance and are associated with subsequent metastases in other organs (Coleman, 2006). These different behaviors may suggest different subtypes of cancers that are yet to be characterized and distinguished. Alternatively, there may be a transition between these phenotypes. In fact, depending on different interaction partners, the same cancer cells may exist in different status in the bone. For instance, while endothelial cells may keep cancer cells in dormancy (Ghajar et al., 2013; Price et al., 2016), osteogenic cells promote their proliferation and progression toward micrometastases (Wang et al., 2015, 2018). Therefore, it is possible that the transition from indolent to aggressive behaviors is underpinned by an alteration of specific microenvironment niches. Detailed analyses of such alteration may be achieved will lead to unprecedented insights into metastatic progression.

Although data presented in this and accompanied studies indicate that cancer cells colonizing the bone acquire intrinsic traits for further dissemination, we cannot rule out systemic effects that may also contribute to this process. At the late stage, bone metastases are known to cause strong systemic abnormality such as cachexia (Waning et al., 2015), which may influence secondary metastasis. Even at early stages before bone metastases stimulate severe symptoms, the disturbance of micrometastases to hematopoietic cell niches may mobilize certain blood cells to migrate to distant organs, which may in turn result in altered metastatic behaviors (Peinado et al., 2017). These possibilities will need to be tested in future research.

## ACKNOWLEDGEMENTS

We would like to thank Zhang Laboratory members and Dr. Lin Tian for helpful input. X.H.-F.Z. is supported by US Department of Defense DAMD W81XWH-16-1-0073 (Era of Hope Scholarship), NCI CA183878, Breast Cancer Research Foundation, and McNair Medical Institute. H.W. is supported in part by US Department of Defense DAMD W81XWH-13-1-0296. We also acknowledge the Pathology Core of Lester and Sue Smith Breast Center, the Genomic and RNA profiling core, the Dan L. Duncan Cancer Center, and Cell Sorting Core (CCSC) at Baylor College of Medicine.

## AUTHOR CONTRIBUTIONS

Conceptualization and Validation, W.Z., I.B., H.W., S.S., H.L., M.N., Z.X., J.L., W.J., J.H., X.H.-F.Z.; Methodology, Formal Analysis and Investigation, W.Z., I.B., H.W., S.S., M.N., Z.X., J.L., W.J., J.H., X.H.-F.Z.; Resources, Y.L., X.H.-F.Z.; Software Data Curation and Visualization, W.Z., S.S., J.H., X.H.-F.Z.; Writing – Original Draft, W.Z., X.H.-F.Z.; Writing – Review & Editing, W.Z., H.L., X.H.-F.Z.; Supervision, X.H.-F.Z.; Project Administration, W.Z., X.H.-F.Z.; Funding Acquisition, H.W., X. H.-F. Z.

---

**Supplementary Figure 1, related to Figure 1.**
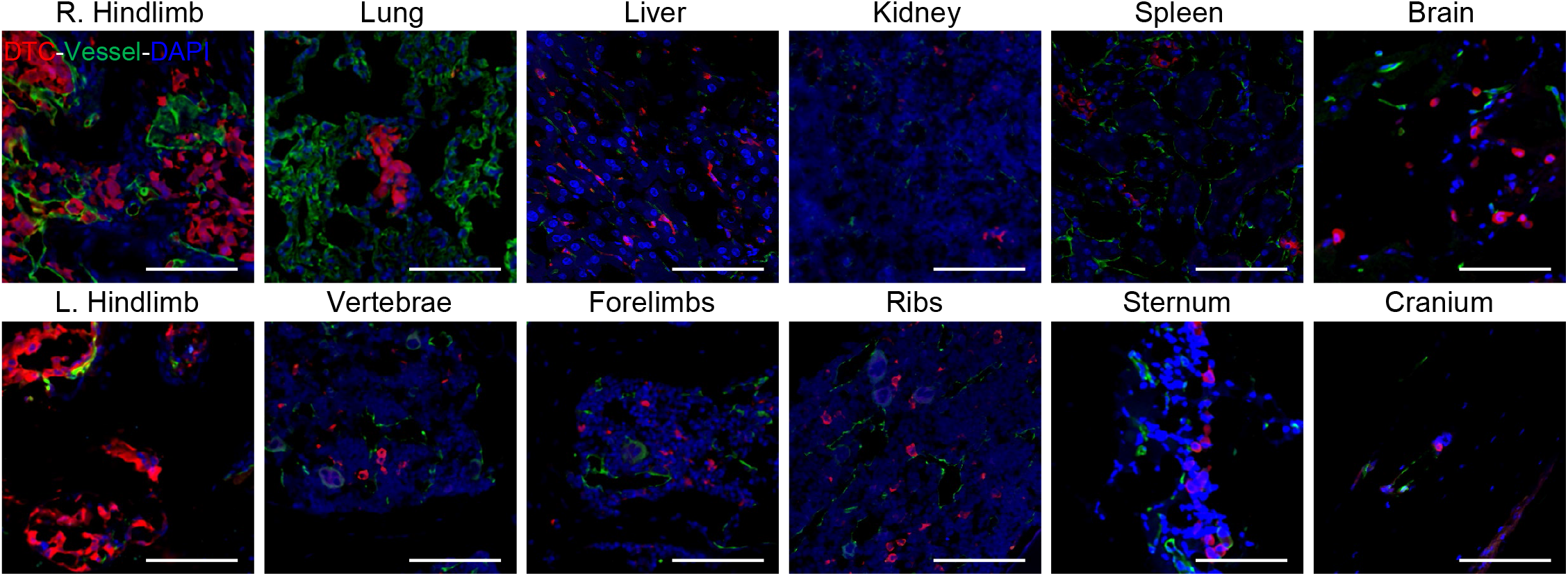
The confocal imaging confirms the presence of fluorescent protein-tagged MDA-MB-231 tumor cells across various tissues. Red, tumor cells; Green, CD31/V-Ecadherin+ endothelium; Blue, nucleus staining. Scale bar, 100 um.

**Supplementary Figure 2, related to Figure 2.**
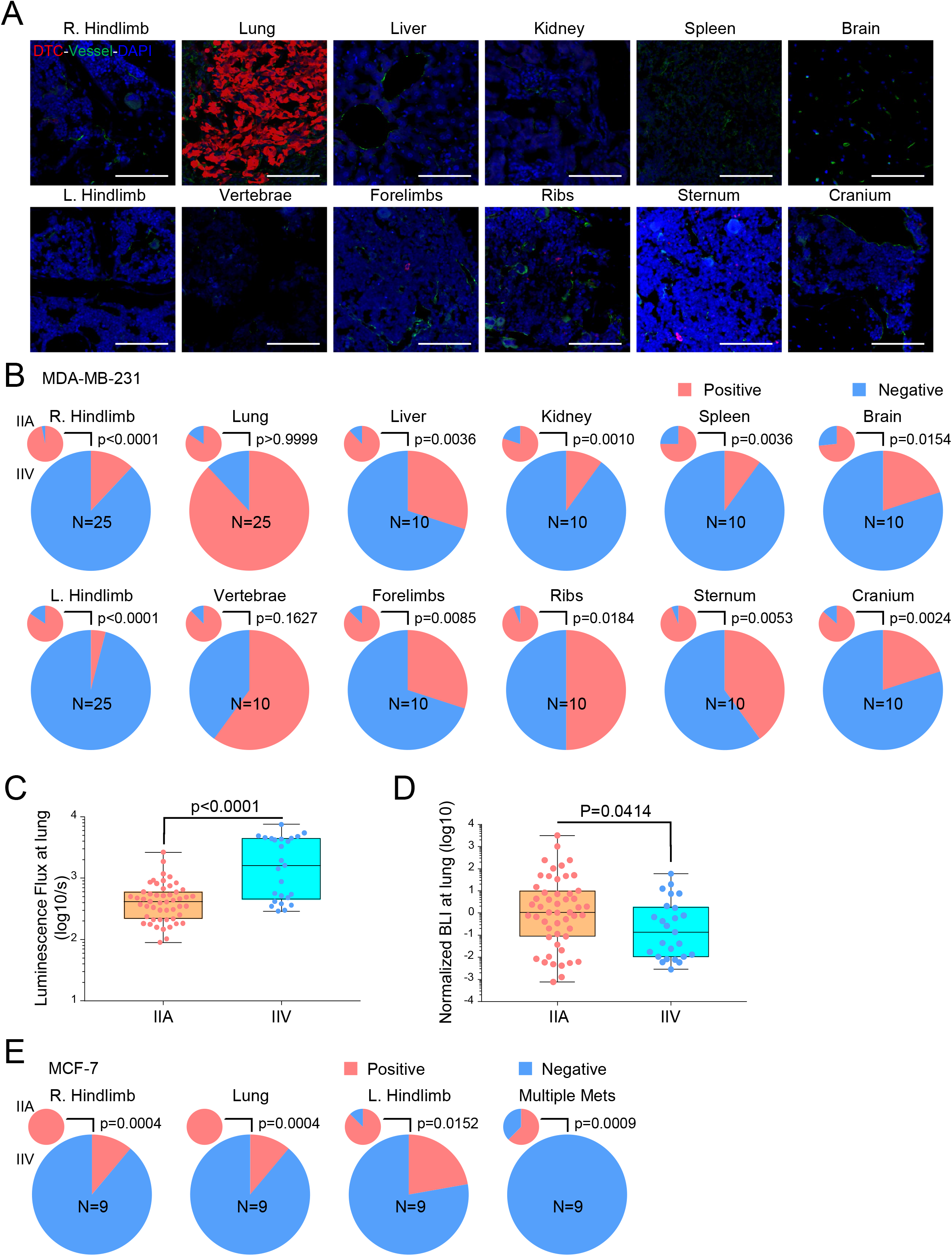
**(A)** Representative confocal images show that less RFP-tagged MDA-MB-231 cells are present in tissues from mice receiving cancer cells through IIV injection. Scale bar, 100 um. **(B)** The frequencies of metastatic involvement across various tissues are significantly decreased in MDA-MB-231 IIV injection model, as compared to those in IIA injection model. **(C)** Comparison of bioluminescence intensity of MDA-MB-231 cells at lung in IIA and IIV injected mice. **(D)** Comparison of the normalized metastatic burden at lung from IIA and IIV injected mice. **(E)** IIA injection of non-metastatic MCF-7 cells in to nude mice led to more frequent metastases in lung and counter-lateral hindlimb (L.Hindlimb), as compared to the IIV model. Overall, no mouse in IIV model generated multiple sites metastases whereas two thirds of mice with IIA injection of MCF-7 cells had metastatic disease in multiple tissues. Fisher’s exact test was used in (B) and (C).

**Supplementary Figure 3, related to Figure 3.**
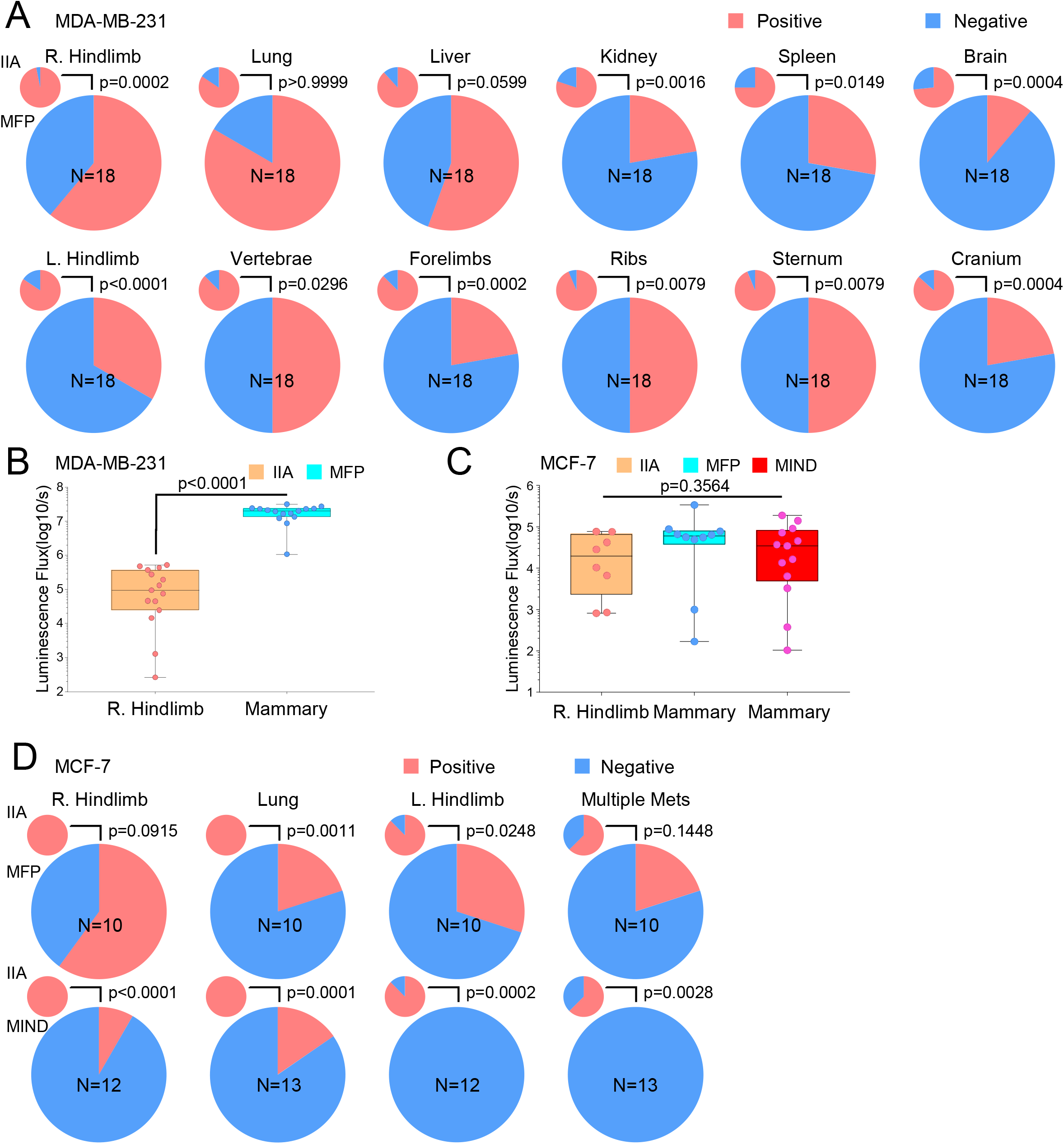
**(A)** The frequencies of metastatic involvement across various tissues in mice with mammary tumors, as compared to those with bone metastases via IIA injection. **(B-C)** Comparison of tumor burden at initial injection sites (bone vs. orthotopic tumors) in different experimental groups. (B), MDA-MB-231 cells, Mann-Whitney test; (C), MCF-7 cells, Kruskal-Wallis test. **(D)** IIA injection of non-metastatic MCF-7 cells in nude mice led to more frequent metastases in lung and counter-lateral hindlimb, as compared to the MFP or MIND model. Rare mice show the multi-organ metastases in both MFP and MIND models, whereas two thirds of mice with bone metastases had metastatic disease in multiple organs. Fisher’s exact test was used in (A) and (D).

**Supplementary Figure 4, related to Figure 4.**
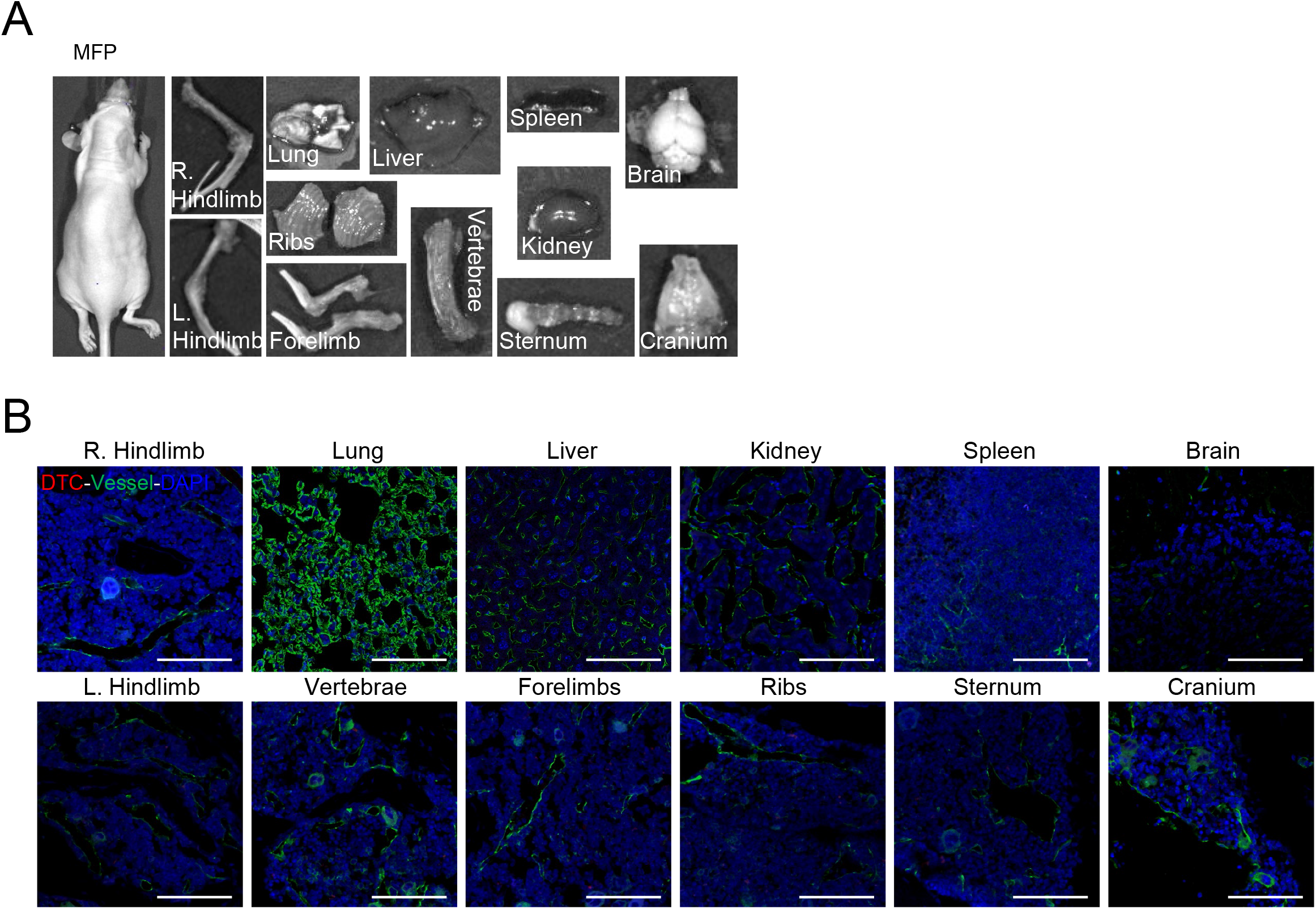
**(A)** Representative *ex vivo* bioluminescent images of multiple organs in parabiotic mice from MFP group, compared to Figure 4E. **(B)** Representative confocal images in search for tumor cells in various tissues of parabiotic mice of MFP group, compared to Figure 4G.

**Supplementary Figure 5, related to Figure 5.**
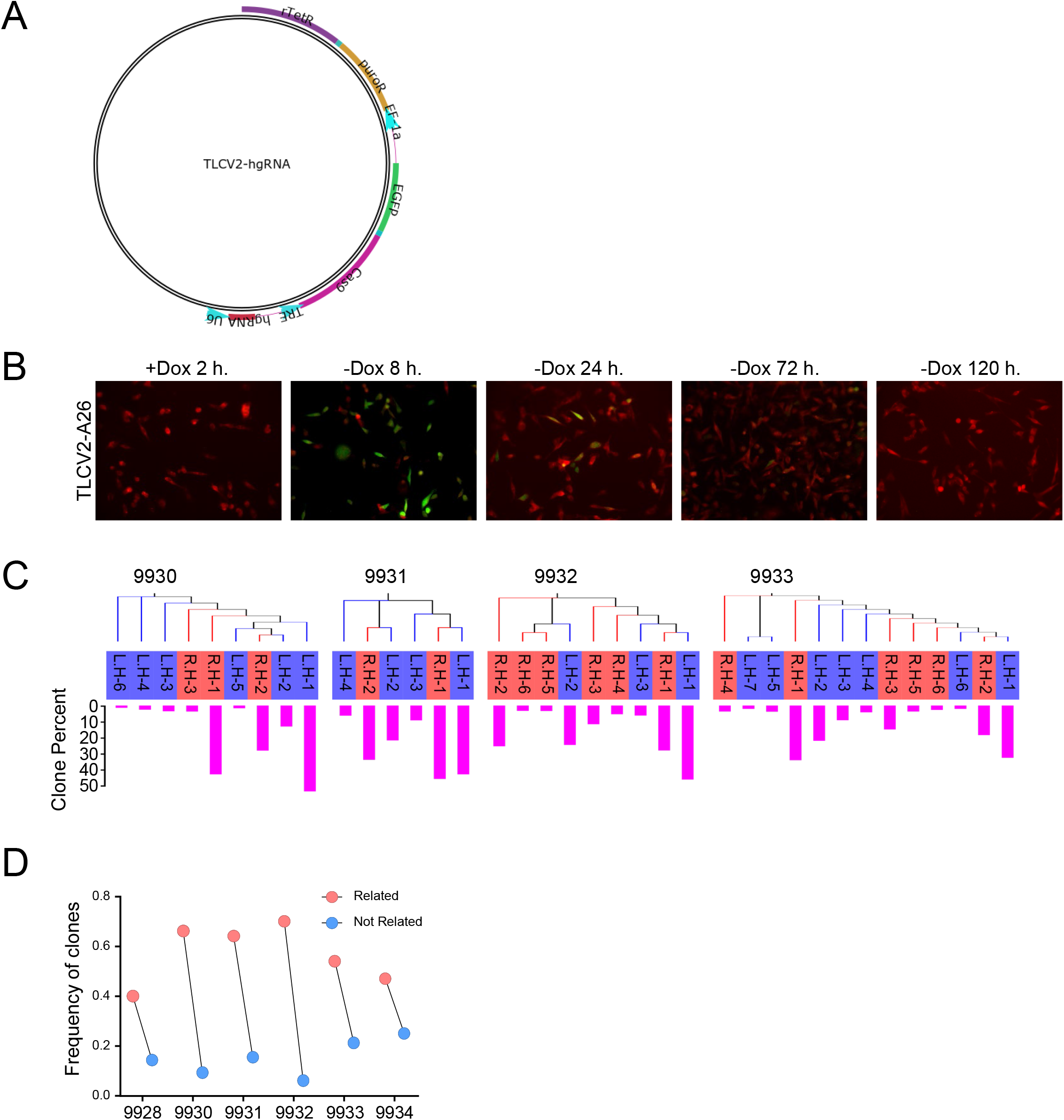
**(A)** The map of TLCV2-hgRNA construct. **(B)** Fluorescence microscopy showing the pulsed expression of Cas9 protein in barcoded MDA-MB-231 cells upon short term of doxycycline treatment. **(C)** Neighbour-joining phylogenetic trees and percentage of clones showing the similarity of barcodes in dominant clones from left hindlimbs and right hindlimbs. The number indicates the different mouse tested. **(D)** The total frequency of related dominant clones in L. Hindlimb, which share the same barcodes with clones from R.Hindlimb, and that of the rest clones were compared.

## STAR Methods

### CONTACT FOR REAGENT AND RESOURCE SHARING

Further information and requests for resources or reagents should be directed to the lead contact Dr. Xiang H.-F. Zhang at

### EXPERIMENTAL MODEL AND SUBJECT DETAILS

#### Cell lines

Human ER-breast cancer cell line MDA-MB-231 and ER+ breast cancer cell line MCF-7, human prostate cancer cell line PC-3, and HEK293T cells were obtained from ATCC. The cells were maintained in DMEM high glucose media supplemented with 10% FBS and 1% penicillin-streptomycin at 5% CO_2_. The SCPs (single cell population) were generated by sorting single parental cells into 96-well plate, and collecting the repopulated clones. MCF-7, MDA-MB-231 cells and respective derivatives were authenticated by MD Anderson Cancer Center CCSG-Characterized Cell Line Core using STR profiling and the profiles of all cell lines matched with NCI public database. Cells were routinely examined for mycoplasma contamination in the lab using PlasmoTest™ Mycoplasma Detection Kit (InvivoGen) and no mycoplasma contamination was detected in the cell lines used in this study.

#### Animals

Athymic nude mice were obtained from Envigo and maintained in the institutional facility for 2 weeks before the experiments. 6- to 8-week-old female or male mice were used for breast cancer or prostate cancer model, respectively, in all *in vivo* studies. In animals injected with MCF-7 and derivative cells, slow-released estradiol tubes were prepared and implanted under the back neck skin of nude mice 1 week prior to the tumor implantation. All animal studies were covered by and conducted in accordance with the protocols approved by the Baylor College of Medicine Institutional Animal Care and Use Committee.

### METHOD DETAILS

#### Inducible homing barcode plasmid

TLCV2 plasmid was a gift from Adam Karpf (Addgene plasmid # 87360) (Barger et al., 2019). The hgRNA oligos, A26-F and A26-R, were synthesized by Integrated DNA Technologies Inc. The oligo sequences are listed in the Key Resources Table. To construct the TLCV2-hgRNA-A26 plasmid, the oligos were annealed and ligated with BsmBI and EcoRI digested TLCV2 plasmid. Plasmid was extracted and the insertion of hgRNA-A26 was confirmed by Sanger-Sequencing.

#### Lentiviral production and transduction

X-tremeGENE HP DNA transfection reagent (Sigma) was used to transfect HEK293T cells with firefly luciferase fused with GFP/mRFP, or TLCV2-hgRNA-A26 together with psPAX2 and pMD2.G packaging plasmids. 48 hours later, the supernatant was harvested and filtered by 0.45 um filter (VWR International). Cells were transduced by the fresh lentivirus with 8ug/ml polybrene (Sigma). Two days later, GFP/mRFP positive cells were sorted by FACS to generate firefly luciferase and GFP/mRFP labelled cell lines. For the barcoding of MDA-MB-231 cells, the transduced cells were selected by 2 ug/ml puromycin for 2 weeks. To induce Cas9 expression *in vitro*, cells were treated with 100ug/ml doxycycline (Sigma) for 2 hours and then rinsed by pre-warmed PBS three times to completely remove doxycycline. For the induction of Cas9 *in vivo*, one dose of 5mg/kg doxycycline was administrated via intra-peritoneal injection weekly.

#### IIA and IIV injection

Both Intra-iliac artery and vein injections were performed using similar procedures as previously described (Wang et al., 2015; Yu et al., 2016). Briefly, animals were anesthetized and restrained on a warming pad. The surgery area was sterilized and a 7-8 mm incision was made between the right hind limb and abdomen to expose the common iliac vessels. 10E4 cells suspended in 100ul PBS were injected by 31G insulin syringe (Becton Dickinson) via iliac artery or vein, respectively. The bleeding was stopped by gently pressing on the incision area with a cotton tip for 5 minutes. Then, the incision was closed by 9 mm wound-clips. The post-operative care was provided and the mice were monitored daily until the removal of wound-clips.

#### Mammary fat pad, tail vein, intra-ductal and intra-cardiac injection

The mammary fat pad injections were performed as described previously (Wang et al., 2015). 10E4 cells mixed 1:1 with growth factor reduced Matrigel Matrix (Corning) were injected to the right fourth mammary gland of mice. For the cross-seeding experiment, mice received the same number of cancer cells mixed with Matrigel at the fourth left mammary gland right after the IIA injection of cancer cells in the right hind limb. For tail vein injection, 2E4 cells in 100ul PBS were injected through tail vein to generate lung metastases comparable to IIA model after 8 weeks. Intra-cardiac injections were performed with 10E4 cells in 100 ul PBS. The intra-ductal injection was performed as previously reported (Nguyen et al., 2000). Briefly, the tip of the fourth nipples was cut off with a sterilized surgical scissor to expose the duct. 10E4 cells were suspended in 30 ul PBS and injected into the nipple duct using 22G blunt needle fitted to a Hamilton syringe.

#### Parabiosis surgery and reverse procedure

The procedure for parabiosis surgery and the subsequent reverse procedure to separate the parabiotic pairs were described previously (Kamran et al., 2013). Briefly, each pair of parabiotic mice were placed in the same cage to ensure harmonious cohabitation two weeks before the experiments. The next week, one mouse within each pair received the tumor implantation via MFP or IIA injection. For parabiosis surgery, both mice were anesthetized by isoflurane and placed back to back on a warming pad. A longitudinal incision starting from the elbow to the knee joints was made and the skin was gently detached from the subcutaneous fascia. The corresponding joints were tightly connected with non-absorbable 4-0 suture. The skin incision was then closed with absorbable 5-0 suture. The parabiotic pairs were closely monitored until full recovery and post-operative care was provided. 7 weeks after the surgery, the reverse procedure was performed to separate the parabiotic pairs.

#### Bioluminescence Imaging and Tissue Collection

Bioluminescence imaging (BLI) of animals was performed weekly using IVIS Lumina II (Perkin Elmer). 100 ul 15 mg/ml D-luciferin (Goldbio) was injected to the anesthetized mice via the retro-orbital venous sinus. If not specified, all the mice were sacrificed 8 weeks after tumor engraftment. For the study of organ distribution of metastases, D-Luciferin was administrated to the live animals before euthanization, and the tissues were dissected sequentially and assessed by *ex vivo* BLI. The regions of interest were manually defined for each type of organs, and the BLI intensities were quantified and presented as the total photon flux/s. Organs showing areas of bioluminescence signal higher than 15 photons/pixel were considered as metastasis positive. The excised tissues were immediately fixed by 4% PFA at 4 °C overnight, cryopreserved with 30% sucrose PBS solution, and then embedded in OCT (Tissue-Tek). For bone tissues, decalcification in 14% PH 7.4 EDTA solution was performed for 1 week before cryopreservation.

#### Immunofluorescent Staining

The frozen sections were prepared by the BCM Breast Center Pathology Core. The immunofluorescent staining was performed with antibodies against mRFP (Rockland, 600-401-379), EGFP (Abcam, 13970), mouse CD31 (R&D Systems, AF3628), and mouse VE-Cadherin (R&D Systems, AF1002). Briefly, the frozen slides were warmed at room temperature for 10 minutes, and rinsed in PBS twice. After 30 minutes penetration with 0.3% Triton X-100 in PBS, the sections were blocked by 10% donkey serum in PBS-GT (2% Gelatin, 0.1% TritonX-100) for I hour at RT. Then the sections were incubated with primary antibodies for overnight at 4°C in a humidified chamber. The next day, the slides were washed and incubated with Alexa Fluor 488 conjugated Donkey anti-Chicken IgY (Jackson ImmunoResearch, 703-546-155), Alexa Fluor 555 conjugated Donkey anti-Rabbit IgG (Thermo Fisher, A31572), and Alexa Fluor 647 conjugated Donkey anti-Goat IgG (Jackson ImmunoResearch, 705-606-147) for 2 hours at RT. The stained sections were then washed and mounted with ProLong^TM^ Gold antifade mountant with DAPI (Thermo Fisher, P36935). Images were acquired by a Zeiss LSM780 confocal microscope.

#### Genomic DNA Extraction from Bone and Lung

The metastatic lesions were excised from mice with *ex vivo* BLI imaging, and the uninvolved tissues were removed. The surgery tools were sterilized with a bead-sterilizer following by 70% isopropanol wash between collections of different lesions. The dissected tissues were snap-frozen in liquid nitrogen and stored in the −80°C freezer until next step. The tissues were first thawed, rinsed with the gDNA lysis buffer from Quick-DNA Miniprep Plus Kit (Zymo Research, D4068), and homogenized with Precellys Lysing Kit (Bertin Instruments, MK28-R). After homogenization, samples were lysed at 55°C for 3 h, followed by 0.33 mg/mL RNase A treatment at 37°C for 15 min. Nucleic acid was further extracted following the manufacturer’s instructions. The final DNA concentration was assessed by NanoDrop 2000 (Thermo Scientific) and human/mouse DNA ratio was examined by q-PCR with primers specifically targeting human and mouse GAPDH genes.

#### Barcode Amplification and Sequencing

Barcodes were amplified by two rounds of PCR. The first round of PCR was performed with 100 ng genomic DNA using Platinum Taq DNA Polymerase (Invitrogen) with Barcode-For and Barcode-Rev primers in 15 cycles. The second round of PCR were performed in a real-time setting and stopped in mid-exponential phase using PowerUp SYBR Green Master Mix (Thermo Fisher) with Barcode-P5-For and Barcode-P7-Rev primers. The sequences of primers are provided in Key Resources Table. PCR products were then column-purified with QIAquick PCR purification Kit (QIAGEN) and assessed with Qubit. The NEBNext Multiplex oligos for Illumina (Dual index primer set 1,NEB, E7600S) and the NEB library preparation kit for Illumina (NEB, E7645S) were used for library preparation as previously described (Kalhor et al., 2017). NextSeq 500/550 High Output Kit v2 (300 cycles) was used and the sequencing was performed by the Genomic and RNA Profiling Core at Baylor College of Medicine.

#### Barcode Data Processing and Analysis

A customized pipeline was used to extract the sequences and counts of barcodes from FASTQ files. Briefly, to identify the barcoding region, each distinct sequence was globally aligned to the A26 reference barcode using pairwise Alignment function of Biostrings package in R. The parameters used for alignment are: 2 for match score, −2 for mismatch score, −10 for gap opening penalty and −0.1 for gap extension penalty. Next, the barcode region from each read was extract, which covers both the spacer and the scaffold sequence of hgRNA. For each sample with the aligned barcodes, the number of reads in each distinct clones was counted and ranked from most frequent to least frequent. The top barcodes that constitute 75% of the total number of reads were considered as dominant clones in each sample. The dominant clones with the sequence at both ends were used for the subsequent analysis of phylogenetic relationship. The sequences and frequencies of these clones in each sample are listed in the supplementary spreadsheet. We used T-Coffee algorithm (European Bioinformatics Institute) to perform the multiple sequence alignment and generate the neighbor-joining phylogenetic trees without distance correction for the dominant clones in each mouse. The visualization of phylogenetic trees and frequencies of clones was achieved by EvolView (He et al., 2016).

#### RNA extraction and qRT-PCR

RNA extraction was performed with Direct-zol RNA miniPrep Kit (Zymo Research). cDNA was generated with RevertAid First Strand cDNA synthesis Kit (Thermo Scientific, K1622) with 1 ug total RNA following the manufacturer’s instructions. Real-time PCR was performed with PowerUp SYBR Green Master Mix (Thermo Fisher) on Biorad CFX Real-Time system. GAPDH mRNA level was used as the internal control to calculate the relative expression level of target genes. The primers are listed in the Key Resources Table.

#### Statistical Test

If not specified, all the data and statistical analysis were generated by GraphPad Prism 7. The types of statistical tests performed were noted in respective legends. P<0.05 in two-tail test was considered as significant.

#### Data availability

The barcoding sequences are provided in the supplementary spreadsheet.

## STAR Methods

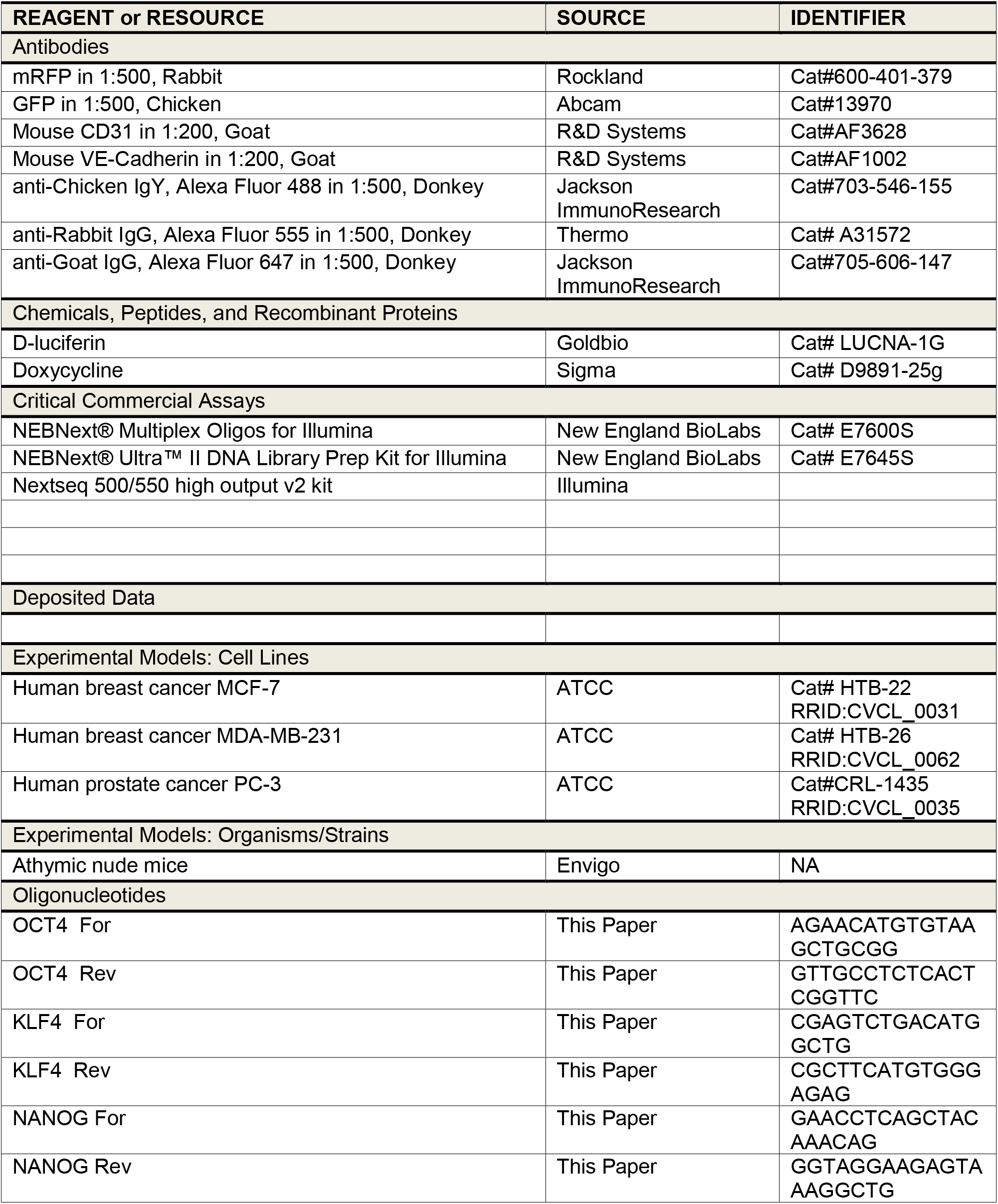
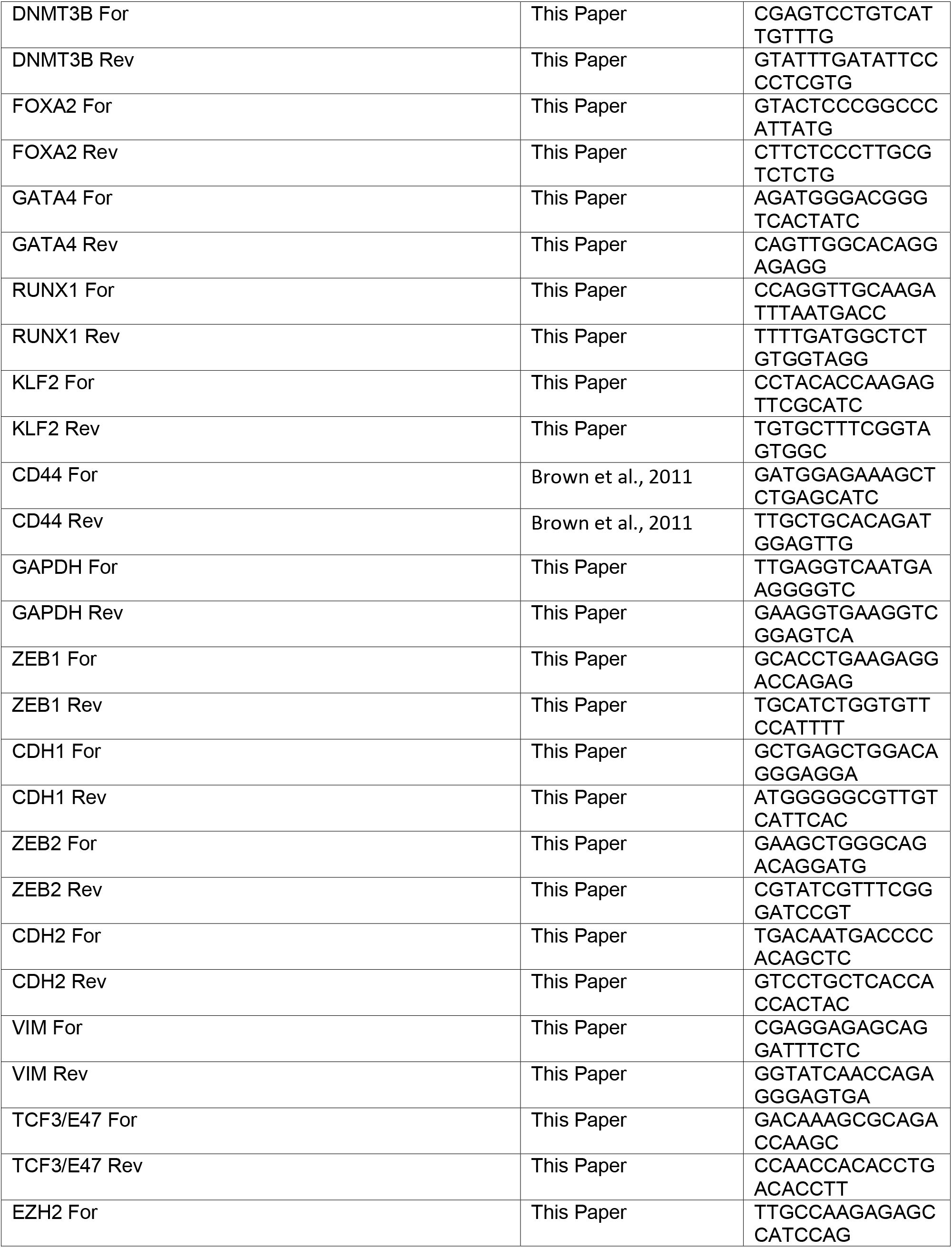
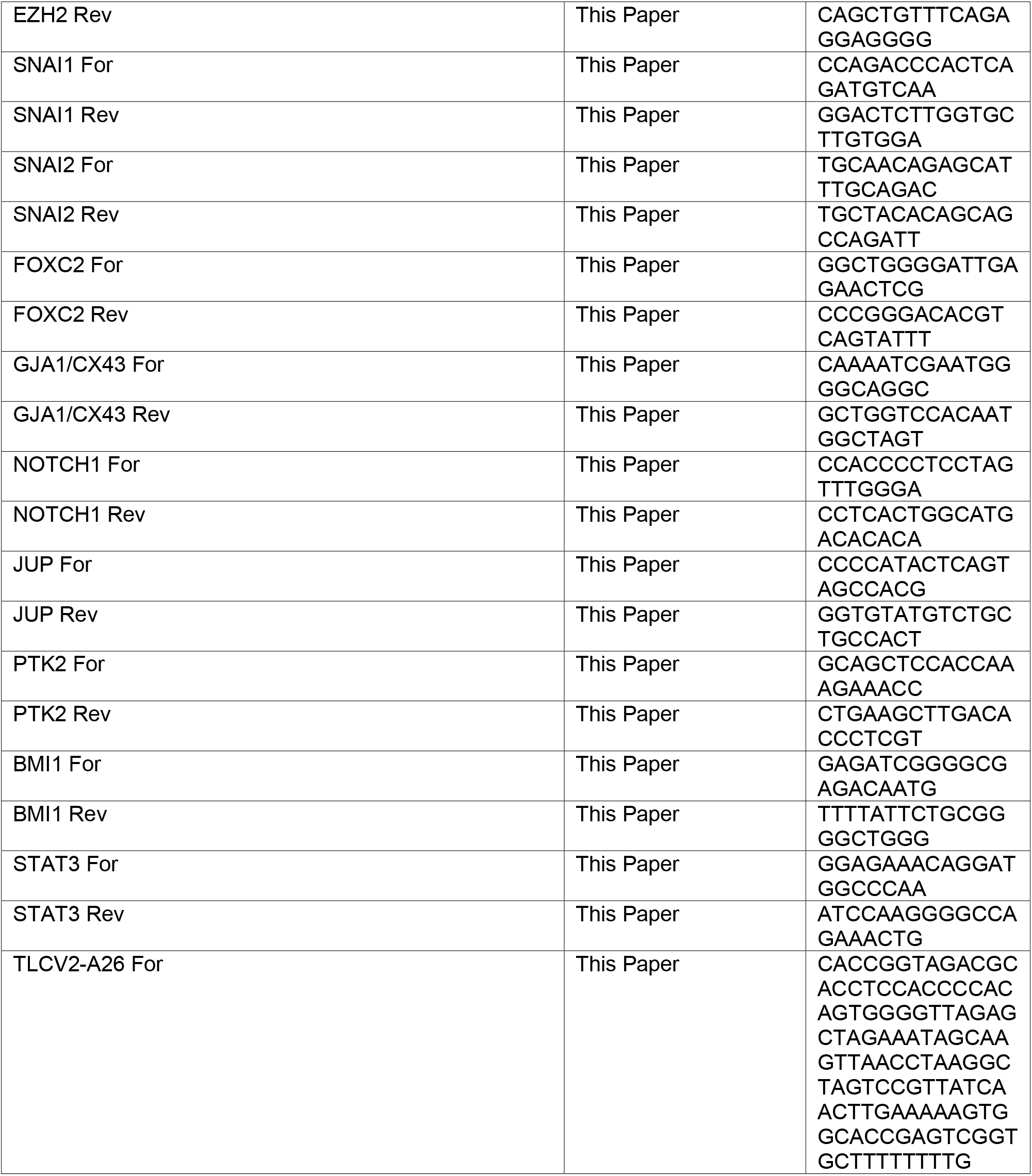
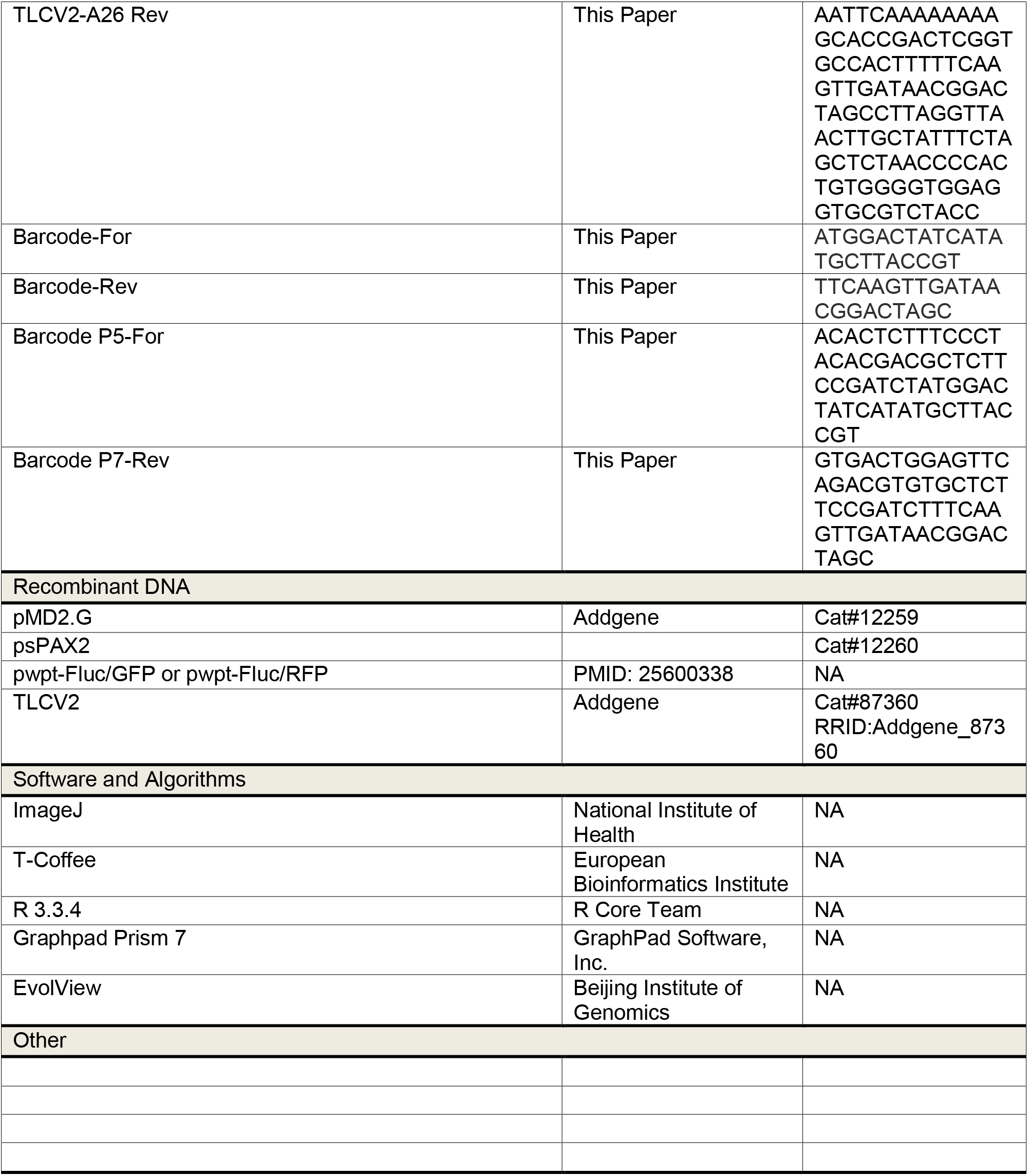
KEY RESOURCES TABLE.

